# Sex-specific axon initial segment plasticity underlies cortical hyperexcitability in trigeminal pain

**DOI:** 10.1101/2025.10.11.681811

**Authors:** Justine Loudan, Karine Herault, Tanguy Damart, Pauline Murail, Alexanne Michon, Sérine Fraiah, François Gabrielli, Xavier Moisset, Radhouane Dallel, Mickael Zbili

## Abstract

Neuropathic pain results from peripheral lesion, causing maladaptive plasticity and central sensitization. Clinical and preclinical studies demonstrate that modifications of primary sensory cortex (S1) activity are essential for neuropathic pain persistence. Rodent studies report heightened S1 pyramidal cell excitability in neuropathic pain model, the origins of which remain debated. The axon initial segment, the action potential trigger zone, is a major determinant of neuronal excitability and is known to undergo structural changes after neural perturbation but its role in chronic pain is poorly understood and no studies have explored its role in cortical hyperexcitability in chronic pain models. Besides, despite a higher prevalence of chronic pain in women, most of preclinical studies have been conducted in males. By integrating electrophysiology, immunohistochemistry, and computational modeling, this study demonstrates that in a trigeminal neuropathic pain rat model, sex-specific structural plasticity of the axon initial segment enhances the excitability of layer 5 pyramidal cells in the somatosensory cortex, potentially driving network-level hyperactivity.

## Introduction

Neuropathic pain, significant public health concern affecting 7–10% of the general population^1^, arises from damage to the peripheral nervous system, leading to maladaptive neural plasticity and central sensitization of pain pathways^2^. Orofacial neuropathic pains constitute a major therapeutic challenge to healthcare professionals^3^. Numerous studies have shown that primary sensory cortex (S1) activity modifications are determinant in neuropathic pain persistence both in human and rodents^4^. Specifically, studies reported an excitability increase of S1 pyramidal cells in neuropathic pain models^5,6^, the origin of which is still debated. Axon initial segment (AIS), the action potential trigger zone, displays structural plasticity following neuronal activity perturbations, causing neuronal excitability modification^7,8^. However, AIS plasticity has been understudied in chronic pain contexts with only two studies reporting an increase of AIS-bearing dorsal root ganglion neurons in a neuropathic pain model^9^ (still under debate^10^) and a distal shift of AIS in spinal dorsal horn inhibitory interneurons in an inflammatory pain model^11^. Therefore, the effect of AIS plasticity at the cortical level in chronic pain is still unknown. Moreover, these two studies, like most of the preclinical pain studies, report results only on male animals. Following Infraorbital Nerve Ligature (IONL), a rodent model of trigeminal neuropathic pain, we found that layer 5 pyramidal cells (L5PCs) of S1 Barrel Field (S1BF) presented hyperexcitability associated with AIS elongation in both sexes. We also found a reduction of soma-AIS distance specifically in females. Using computational approaches, we showed that AIS plasticity underlies L5PCs hyperexcitability in neuropathic animals leading to neuronal network hyperactivity.

## Materials and methods

All details are provided in Supplemental material.

### Animals

Experimental protocols on female and male Sprague–Dawley rats were approved by the Committee of Animal Research at the University of Clermont Auvergne, France and in accordance with the International Association for the Study of Pain.

### Surgery

Infraorbital Nerve Ligation (IONL) was performed following a previously established surgical procedure^12^.

### Experimental protocols

Allodynia score was obtained by application of innocuous Von Frey filament in the infraorbital nerve territory ipsilaterally to the surgery^13^. Patch-clamp recordings of L5PCs were made on acute slices of S1BF from the contralateral side of the surgery. AIS and soma morphological measurements were obtained by co-immunostaining of patch-clamp slices using anti-CamKIIα and anti-ankyrinG antibodies.

### Computational modelling

Hodgkin-Huxley models were done in NEURON 8.2.6 software using a ball-and-stick morphology composed of one soma, one hillock and one AIS which morphologies were modified in the different tested conditions. The network modelling was performed using python self-written codes, employing conductance-based adaptative leaky integrate-and-fire (CAd-LIF) neurons, whose parameters were determined to fit excitability curves of recorded neurons.

### Statistics

Statistical analyses for groups comparison were performed using mixed linear model regressions (MLMR)^14^ followed by pairwise t-tests based on marginal means and variances derived from the models. Correlation parameters with day post-surgery were performed using Pearson’s linear regressions test. All the analyses were made using statsmodels and scipy python libraries.

## Results

### Trigeminal nerve injury induces mechanical allodynia

Assessing the facial allodynia score^12,13^ by application of an innocuous Von Frey filament at the vibrissae pad level, we found that infraorbital nerve ligation (IONL), but not Sham procedure, produced a severe ipsilateral face allodynia from the 10^th^ day post-surgery both in male (Fig. 1A) and female rats (Fig. 1B). However, we did not find difference between allodynia scores in males and females at any of the tested days suggesting that IONL induced a similar allodynia in both sexes.

**Figure 1.**
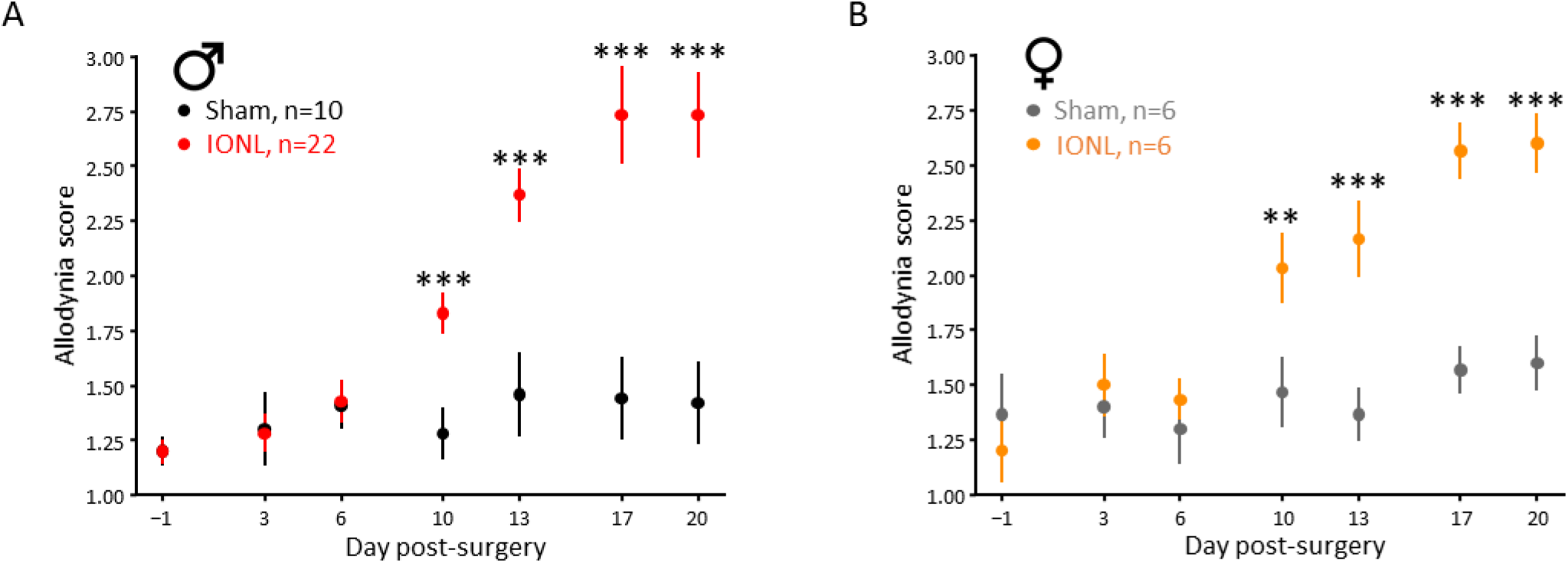
Infra-orbital Nerve Ligation induces similar mechanical allodynia in both sexes. **(A)** Allodynia scores induced by non-nociceptive Von-Frey filament application at vibrissae pad zone in male rats. Note that allodynia score does not increase following a Sham surgery while rats begin to show facial allodynia from the 10th day after IONL surgery (Sham animals: black, *n =* 10; IONL animals: red, *n =* 22). **(B)** Allodynia scores induced by non-nociceptive Von-Frey filament application at vibrissae pad zone in female rats. Note that allodynia score does not increase following a Sham surgery while rats begin to show facial allodynia from the 10th day after IONL (Sham animals: grey, *n =* 6; IONL animals: orange, *n =* 6).

### Hyperexcitability of layer 5 pyramidal cells in neuropathic condition

S1 hyperexcitability is one of the sources of neuropathic pain^5,6^. To determine the cellular origin of this hyperexcitability, we performed patch-clamp recordings of layer 5 pyramidal cells (L5PCs) in S1BF contralateral to the surgery. We found an increase of L5PCs firing frequency in IONL animals both in males (Fig. 2A) and females (Fig. 2B) with input/frequency (I/F) curve gain increase in both sexes (Fig. 2D) and rheobase decrease only in males (Fig. 2C). Importantly, while the neurons recordings were acquired from 11 to 42 days post-surgery (d.p.s), we found no correlation between d.p.s and rheobase or I/F curve gain under any conditions (Supplementary Fig. 1A-B), consistent with previous results showing IONL-induced allodynia from 12 to at least 120 d.p.s^13^. Interestingly, independently of the condition, L5PCs displayed a larger excitability in females than in males (Fig. 2C-D, Supplementary Fig. 1C-D). In conclusion, IONL induced an L5PCs excitability increase on the contralateral S1BF in both sexes.

**Figure 2.**
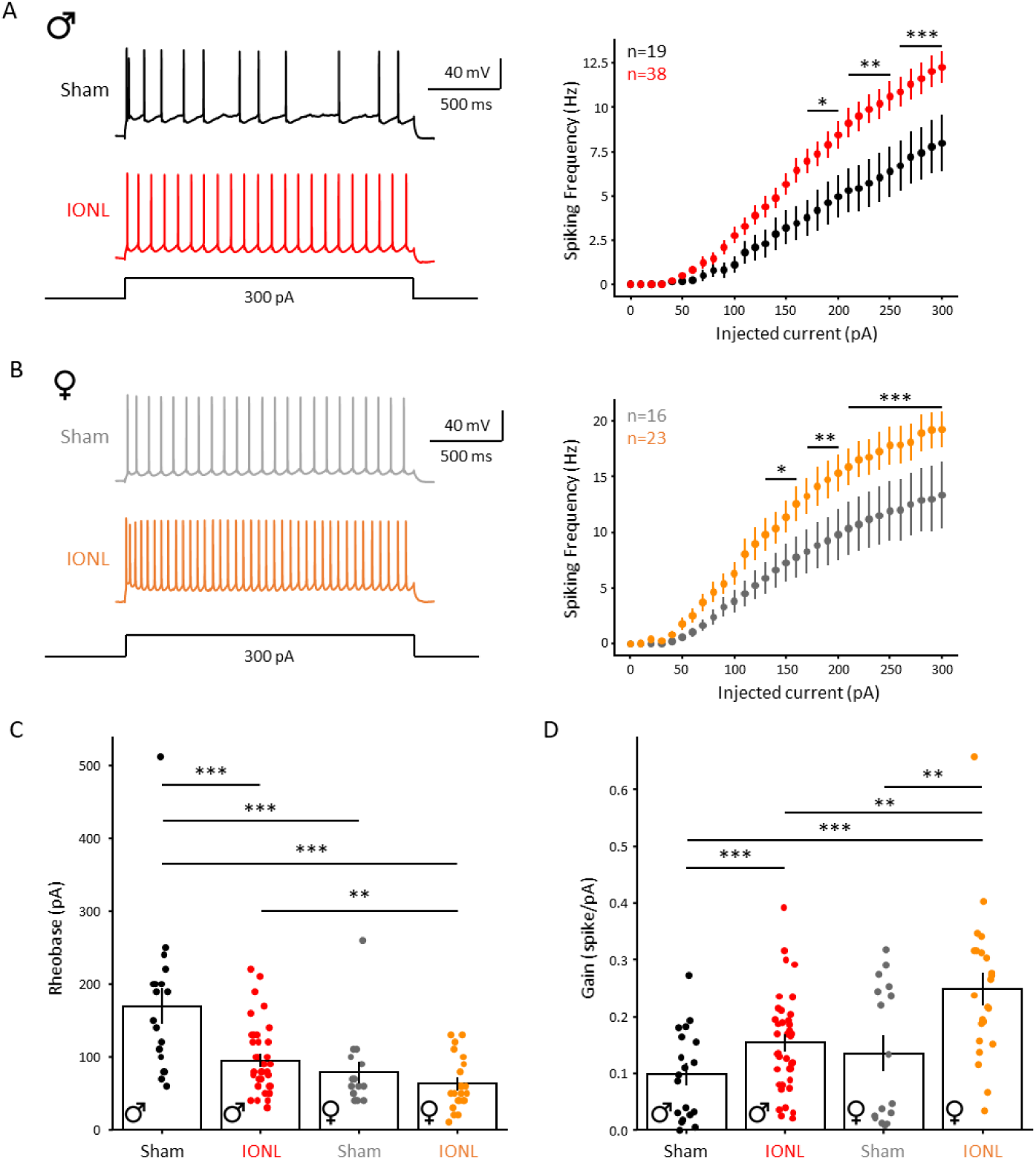
Increase of Barrel cortex Layer 5 pyramidal cells excitability following IONL. **(A)** Increase in L5PC excitability in IONL animals (red) compared to Sham animals (black) in male rats. Left, Example current-clamp traces recorded in L5PC in Sham (black) and in IONL (red) animals. Right, Average input/output curves. L5PCs display a larger excitability in IONL (red; *n =* 38 cells in 14 animals) than in Sham animals (black; *n =* 19 cells in 11 animals). **(B)** Increase in L5PC excitability in IONL animals (grey) compared to Sham animals (orange) in female rats. Left, Example current-clamp traces recorded in L5PC in Sham (grey) and in IONL (orange) animals. Right, Average input/output curves. L5PCs display a larger excitability in IONL (orange; *n =* 23 cells in 7 animals) than in Sham animals (grey; *n =* 16 cells in 6 animals). **(C)** L5PC rheobase in Sham and IONL male and female rats. Note that rheobase is reduced following IONL in male but not in female rats. Note that female L5PC display smaller rheobase than male in Sham animals, suggesting a sexual dimorphism of excitability for this neuronal type. **(D)** L5PC I/F curve gain in Sham and IONL male and female rats. Note that gain is increased following IONL in male and in female rats.

Analyzing L5PCs electrophysiological parameters, we found that IONL induced an increase of membrane resistance specifically in males (Supplementary Fig. 2A) but no modification of membrane capacitance, resting membrane potential, AP threshold, AP half-width or AP peak (Supplementary Fig. 2B-F). No correlation between the d.p.s and most electrophysiological parameters was found, except for a depolarization of the resting membrane potential in female Sham animals (Supplementary Fig. 3). Interestingly, L5PCs displayed a smaller membrane capacitance and a larger membrane resistance in females than in males (Supplementary Fig. 2A-B), consistent with the larger excitability found in females. In conclusion, IONL induced a L5PCs excitability increase on the contralateral S1BF in both sexes without altering electrophysiological parameters, except for an increase of membrane resistance in male animals.

### Axon initial segment plasticity of layer 5 pyramidal cells in neuropathic condition

Axon initial segment (AIS) is a major determinant of cortical neurons excitability. Numerous studies have shown that networks activity perturbations lead to AIS length and soma-AIS distance plasticities causing neuronal excitability modifications^7,8^. Moreover, peripheral injury can induce S1 activity pertubations leading to neuronal hyperexcitability^5^. Therefore, we asked if S1BF L5PCs hyperexcitability in IONL animals resulted from AIS plasticity. We performed co-immunostaining of CamKIIα, a pyramidal cells marker^15^, and Ankyrin G, an AIS marker^7^, on S1BF slices. We found that IONL induced a AIS length increase of CamKIIα^+^ neurons both in males and females (Fig. 3C) and a reduction of soma-AIS distance in specifically females (Fig. 3D). We did not find correlation between the d.p.s and AIS parameters, except for a soma-AIS distance reduction in Sham males (Supplementary Fig. 4A-B). In conclusion, IONL induced L5PCs AIS elongation on contralateral S1BF in both sexes, and reduced soma-AIS distance only in females.

**Figure 3.**
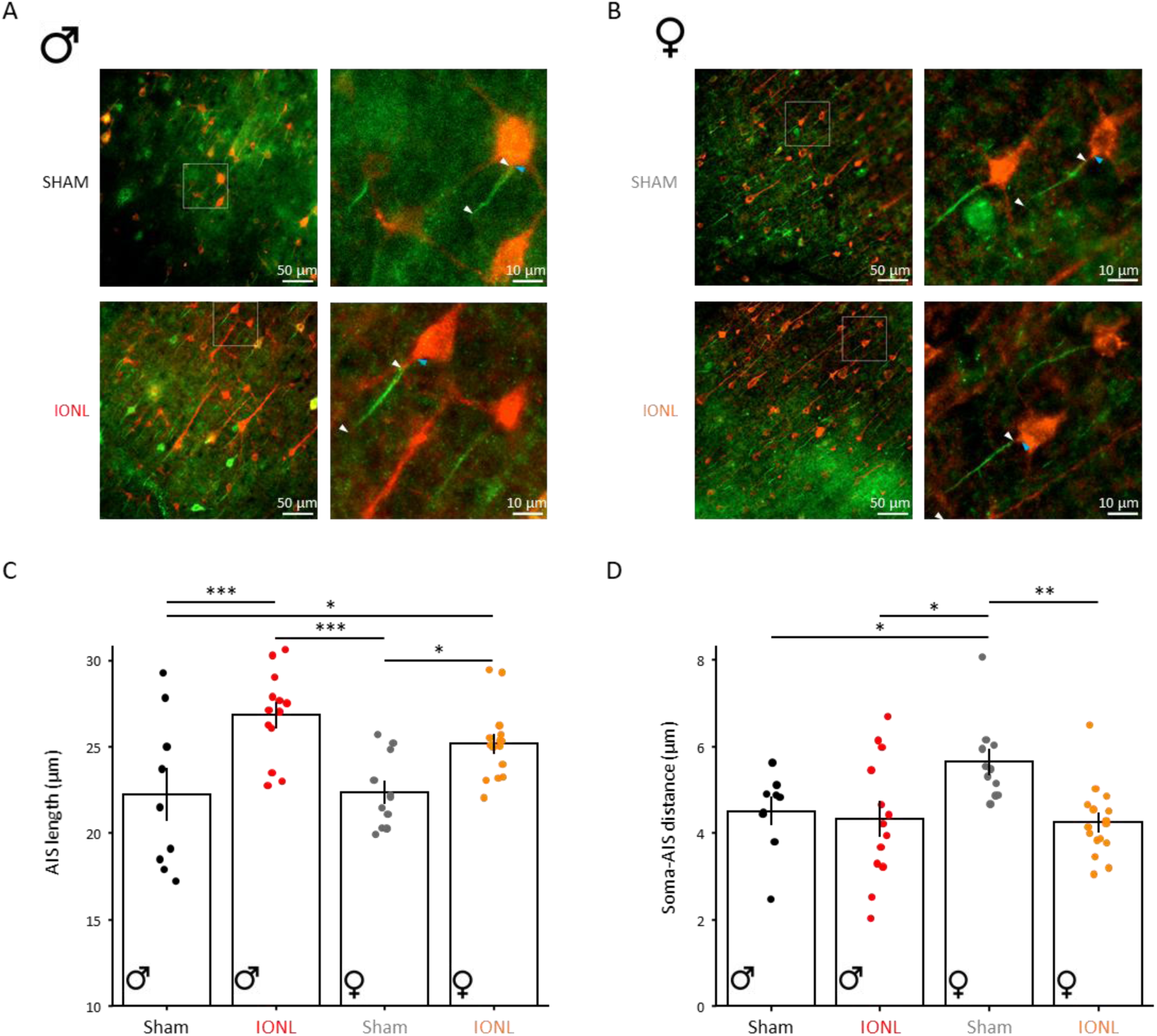
Axon initial segment plasticity in Layer 5 pyramidal cells following IONL. **(A)** Left, Layer 5 barrel cortex region of Sham (upper panel) and IONL (lower panel) male rats slices stained with CamKIIα (red) and Ankyrin G (green) antibodies. Right, Amplified images of staining in the regions indicated in the left panels (white boxes). **(B)** Left, Layer 5 barrel cortex region of Sham (upper panel) and IONL (lower panel) female rats slices stained with CamKIIα (red) and Ankyrin G (green) antibodies. Right, Amplified images of staining in the regions indicated in the left panels (white boxes). **(C)** L5PC AIS length in Sham and IONL male and female rats (Sham male : 799 AIS from 9 slices in 4 animals, IONL male : 1032 AIS from 13 slices in 6 animals, Sham female : 937 AIS from 11 slices in 5 animals, IONL female : 1763 AIS from 16 slices in 6 animals). Note that AIS length is increased following IONL both in male and in female rats (each dot represents the mean of slice). No difference in AIS length between male and female was found in Sham condition nor in IONL condition. **(D)** L5PC soma-AIS distance in Sham and IONL male and female rats (each dot represents the mean of slice). Note that soma-AIS distance is decreased following IONL in female but not in male rats. Note that female L5PC display larger soma-AIS distance than male L5PC in Sham animals, suggesting a sexual dimorphism of this parameter for this neuronal type.

### Layer 5 pyramidal cells displayed smaller soma surface in females than males

Patch-clamp recordings showed that L5PCs displayed a smaller membrane capacitance in females than in males (Supplementary Fig. 2) that could result from a sexual dimorphism in soma size^16^. Moreover, somato-dendritic compartment decrease have been reported in inflammatory pain^17^. Therefore, using CamKIIα stainings, we measured soma surface in Sham and IONL animals. While we found no effect of IONL on soma surface, CamKIIα^+^ neurons displayed a smaller soma surface in females than in males both independently of the condition (Supplementary Fig. 4C). No correlation was found with d.p.s (Supplementary Fig. 4D). In conclusion, we unraveled a sexual dimorphism in S1BF L5PC morphology with smaller soma area in females than in males.

### Axon initial segment plasticity is sufficient to produce IONL-induced increase of excitability

Our results showed a co-occurrence of excitability increase and AIS plasticity in S1BF L5PCs following IONL. While it is widely assume that AIS elongation induces excitability increase^7^, soma-AIS variations effect distance is debated^18^. Importantly, computational studies have focused on AIS plasticity effect on spike initiation without exploring the full I/F curve^18,19^. To determine if the IONL induced-AIS plasticity can explain the L5PCs hyperexcitability, we built a ball-and-stick Hodgkin-Huxley model composed of a soma, a hillock and an AIS (Supplementary methods). For the AIS length and soma-AIS distance, we averaged the experimental values of the conditions showing no differences (Fig. 3C-D). Therefore, AIS length was 22.3 µm in Sham and 26 µm in IONL condition in both sexes. Soma-AIS distance was 5.6 µm in Sham female condition and 4.4 µm in other conditions (Supplementary Table 2). Using multi-objective optimization^20^, we built the male model by fitting the I/F curve observed in Sham males (Fig. 4A, black). In this model, elongating the AIS length up to IONL value reproduced the IONL-induced excitability increase (Fig. 4A, red). The female model was built by a simple reduction of the soma diameter. In this model, AIS elongation and soma-AIS distance reduction was enough to reproduce the IONL-induced excitability increase (Fig. 4C). Therefore, AIS plasticity is sufficient to explain excitability modifications in neuropathic animals of both sexes. Moreover, varying the AIS length in a large range of values had a strong impact on excitability while soma-AIS distance had a much smaller effect (Supplementary Fig. 5). Therefore, our model suggests that AIS length is a major determinant of L5PCs excitability and that AIS plasticity is at the origin of excitability increase in IONL animals.

**Figure 4.**
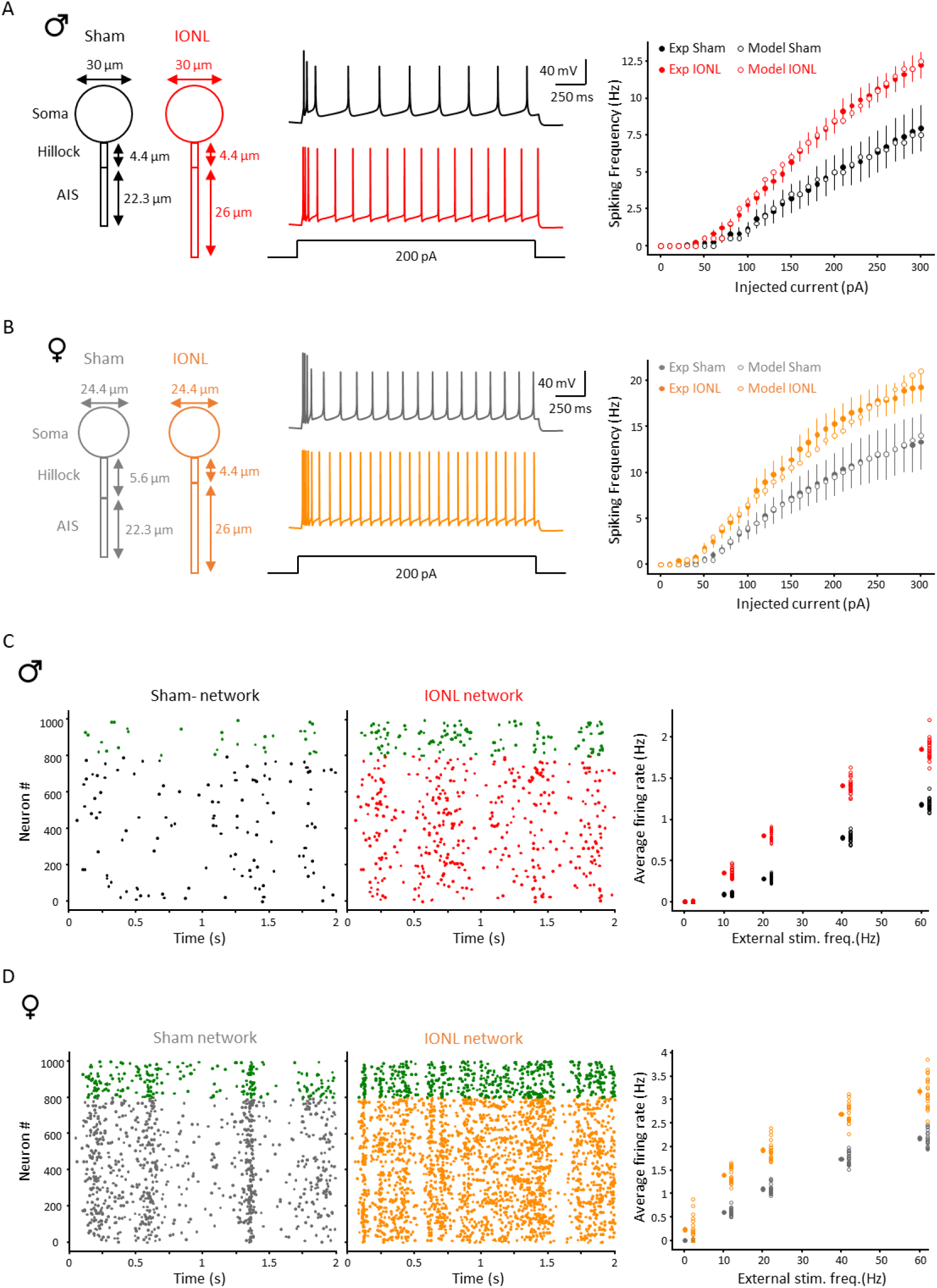
Computational modelling: IONL excitability increase is explained by AIS plasticity and induces hyperactivity at the network level. **(A)** Hodgkin-Huxley single neuron model: AIS plasticity displayed by IONL male rats is sufficient to reproduce excitability increase experimentally observed. **(B)** Hodgkin-Huxley single neuron model: AIS plasticity displayed by IONL female rats is sufficient to reproduce excitability increase experimentally observed. **(C)** Network model for male rats: IONL induced excitatory neurons excitability increase produces network hyperactivity. **(D)** Network model for female rats: IONL induced excitatory neurons excitability increase produces network hyperactivity.

### Excitability increase of layer 5 pyramidal cells induces network hyperactivity in a computational model

In order to observe L5PCs hyperexcitability effect on network behavior, we used a computational approach. Network models were composed of 1000 conductance-based adaptative leaky integrate-and-fire (CAd-LIF) neurons^21^ (80% pyramidal cells and 20% Fast-spiker GABAergic interneurons). Four different CAd-LIF models reproduced L5PCs excitability in Sham and IONL conditions in both sexes (Supplementary Fig. 6A-B) and one CAd-LIF model reproduced Fast Spiker (FS) interneurons excitability from recordings obtained in all conditions (*n=*9 cells, Supplementary Fig. 6C). We built a Sham male network composed of 800 Sham male pyramidal cells and 200 FS interneurons with external inputs simulated by stimulation of each pyramidal cell with a different 10Hz train of excitatory synapses (Fig. 4C, black rasterplot). Replacing Sham male pyramidal cells by IONL male pyramidal cells in this network leaded to a steep increase of the network activity (Fig. 4C, red rasterplot). This IONL-network hyperactivity was found in a large range of external input frequencies (Fig. 4C, right). Building a female network using the same approach, we also found larger network activity in IONL conditions (Fig. 4D). In conclusion, network modelling predicts that AIS plasticity-induced pyramidal hyperexcitability leads to cortical network hyperactivity in neuropathic conditions.

## Discussion

The present study investigated the cellular mechanisms underlying neuropathic pain-associated hyperexcitability in S1BF following infraorbital nerve ligation (IONL) both in male and female rats. Our results demonstrate that IONL-induced robust ipsilateral facial allodynia is associated with a significant increase in the AIS length and intrinsic excitability of layer 5 pyramidal cells (L5PCs) in the contralateral S1BF. We found notable sex differences in L5PCs electrophysiological properties, soma size and AIS plasticity.

### IONL-induced Allodynia

Consistent with previous studies, IONL produced severe and persistent ipsilateral mechanical allodynia^12,13^. While some studies have integrated female in their experiences^22^, our study is the first one displaying a quantitative comparison of sexes in IONL model, showing no difference in the allodynia development with this paradigm.

### Hyperexcitability of L5PCs in chronic pain

The increase in L5PCs excitability observed in IONL animals is in line with the concept of central sensitization, where peripheral nerve injury leads to hyperexcitability in central pain-processing regions^2^. Increase of S1 pyramidal cells excitability in neuropathic pain context have been previously reported^5,6^ but our study report for the first time a similar excitability increase in both sexes for a trigeminal neuropathic pain. Moreover, we found that, regardless of the conditions, S1BF L5PCs display greater excitability, greater membrane resistance and smaller capacitance in females than in males most likely attributable to the smaller soma size found in females. Greater excitability of pyramidal cells in females have been found in prelimbic cortex^23^ but not prefrontal or insular cortices^24,25^. In rats, smaller soma size in females has been found in nucleus accumbens but not in other striatal regions^26^ and has not been found in human cortex^27^. Hence, the sexual dimorphism in electrophysiology and soma size we found may arise from the specific region we studied. Therefore, further efforts are needed to explore sexual dimorphism at the neuronal level, particularly to refine our understanding of the mechanisms leading to chronic pain behavior.

### AIS Plasticity of L5PCs in chronic pain

The AIS is a critical regulator of neuronal excitability, and its plasticity has been implicated in various forms of neuronal adaptation and pathology^7^. However, AIS plasticity has been largely unstudied in chronic pain contexts. In fact, to this date, only two recent studies reported AIS plasticities in pain models : a debated^10^ increase of AIS-bearing DRG neurons in neuropathic pain^9^ and distal shift of AIS in spinal dorsal horn inhibitory interneurons in inflammatory pain^11^. Our results show that IONL induces an elongation of S1BF L5PCs AIS in both sexes, and a reduction of soma-AIS distance specifically in females. Moreover, using Hodgkin-Huxley modeling, we showed that AIS plasticity is enough to explain IONL-induced L5PC hyperexcitability both in males and females. Notably, in line with the sexual dimorphism in soma size we found, our results showed that the same model can fit male and female excitability curve by simply modifying soma diameter. Therefore, modeling suggests that AIS plasticity is a major determinant of chronic pain associated S1 modifications and that somatic morphological dimorphism is at the origin of the greater pyramidal excitability observed in females.

### Network-Level Consequences of Hyperexcitability

At the network level, our simulations predict that the IONL-induced L5PCs hyperexcitability leads to cortical hyperactivity, a hallmark of central sensitization in chronic pain^2^. The network modeling suggests a stronger hyperactivity in females than in males but this arises from the fact that males and females networks models are the same except from the L5PCs excitability which we found to be greater in females. Therefore, next studies will have to unravel potential sexual dimorphism in synaptic connections or GABAergic interneurons excitability to confirm or disprove this prediction.

### Neuropathic pain and homeostatic plasticity

Several studies suggest that maladaptative homeostatic plasticity is a core mechanism for the central networks hyperactivity leading to neuropathic pain^5,28^. Here, we found that hyperexcitability in S1 arises in part from AIS plasticity, a mechanism implicated in homeostatic plasticity^7^. Therefore, it will be of great interest to understand in future studies if AIS plasticity found in neuropathic conditions arises from homeostatic plasticity in order to reverse this process via cortical stimulations^5^ or pharmacology treatments^29^.

In conclusion, our findings contribute to the growing body of literature on cortical plasticity in chronic pain and the sex-specific mechanism we unravaled highlight the importance of considering sex as a biological variable in chronic pain research^30^.

## Abbreviations

AIS: axon initial segment
AnkG: AnkyrinG
CaMKIIα: Calcium/calmodulin-dependent protein kinase type II subunit alpha
IONL: infraorbital nerve ligation
S1BF: primary cortex barrel field
L5PC: layer 5 pyramidal cells
FS: Fast-Spiker
d.p.s: days post-surgery
CAd-LIF: conductance-based adaptative leaky integrate-and-fire

## Data availability

All information necessary to evaluate the findings of the paper are included in the paper. All the codes and .csv dataset files used are available at https://github.com/mizbili/Axon-initial-segment-and-facial-neuropathy. Additional data can be provided from the authors on request.

## Acknowledgements

Author contributions: J.L., R.D. and M.Z. conceived the project. J.L, K.H, P.M, A.M, S.F and M.Z. performed the experiments. J.L., F.G. and M.Z. analyzed the data. T.D and M.Z., performed modeling. J.L., X.M., R.D. and M.Z. wrote the manuscript. M.Z. supervised the project.

## Supplementary material

### Materials and methods

#### Animals

Young adult female and male Sprague–Dawley rats (125–150 g) were obtained from Charles River and housed 3-4 per cage in a controlled environment (inverted light/dark cycles, lights on at 07:00 p.m., 22 °C, food and water ad libitum) for at least 4 days before the beginning of the experiments. Animal protocols were approved by the Committee of Animal Research at the University of Clermont Auvergne, France, and authorized by the French Superior Education and Research Ministry (Authorization no. 50943). Protocols were in accordance with the International Association for the Study of Pain (https://www.iasp-pain.org/resources/guidelines/iasp-guidelines-for-the-use-of-animals-in-research/), the European Directive (2010/63/EU) and ARRIVE guidelines.

#### Surgery

Infraorbital Nerve Ligation (IONL) was performed following a previously established surgical procedure^1–3^. Briefly, after animals were anesthetized using ketamine (50 mg/kg)/xylazine (10 mg/kg), the right Infraorbital nerve was exposed just caudal to the vibrissal pad and two silk ligatures (Peters surgical 4/0 ref 2616) were loosely tied around the nerve just cranial to its exit from the Infraorbital foramen. The ligatures were separated by a 2-3 mm interval. Skin incision was closed with two sutures points (4/0 nylon). For Sham procedure, the right Infraorbital nerve was similarly exposed but not tied.

#### Behavioral testing

Habituation sessions were performed during 5 days before pre-operative testing in order to train rats to not respond to application of non-nociceptive Von Frey filament (VF-6 g or VF-8g determined-before surgery for each animal) in the Infraorbital Nerve (ION) territory (skin in the vibrissae zone). Mechanical stimulation testing was performed on pre-operative day −1 and on post-operative days, +3 and+6, +10, +13, +17 and +20. For the tests sessions, the non-nociceptive VF filament was applied in the ION territory and a allodynia score was determined: (score 0) no response (score 1) detectio*n=*the rat turns the head toward the stimulating object and the stimulus object is then explored (score 2) withdrawal reactio*n=*the rat turns the head slowly away or pulls it briskly backward when the stimulation is applied (score 3) escape/attack=the rat avoids further contact with the stimulus object, or actively by attacks the stimulus object (score 4) freezing^1,3^. The average allodynia score of 5 VF-6g applications was obtained for each rat per test session.

#### Patch-clamp recordings

Recordings of layer 5 pyramidal neurons were obtained using ex-vivo patch-clamp recordings on acute slices of S1BF (primary somatosensory cortex Barrel Field) from the contralateral side of the surgery. The recordings were made on Sham or IONL animals after the mechanical allodynia had developed (Sham: day post-surgery: 11 to 38, mean: 24.4 ± 1.1; IONL: 14 to 42, mean: 26.2 ± 1.1). Firstly, rats were deeply anesthetized with ketamine (50 mg/kg)/xylazine (10 mg/kg) and ice-cold perfused with the slicing solution (92 mM n-methyl-d-glutamine, 30 mM NaHCO3, 25 mM d-glucose, 10 mM MgCl2, 2.5 mM KCl, 0.5 mM CaCl2, 1.2 mM NaH2PO4, 20 mM Hepes, 5 mM sodium ascorbate, 2 mM thiourea, and 3 mM sodium pyruvate, bubbled with 95% O2–5% CO2 (pH 7.4)) and killed by decapitation. After dissection, S1BF slices were cut in the ice-cold slicing solution using a vibratome (Leica VT1200S). Before recordings, slices recovered 30 minutes at 35°C and 30 minutes at room temperature in artificial cerebrospinal fluid (aCSF) (128 mM NaCl, 3 mM KCl, 2.5 mM CaCl2, 1.5 mM MgSO4, 0.6 mM NaH2PO4, 25 mM NaHCO3, 10 mM glucose (300-310 mOsm.kg-1), pH 7.4, bubbled with 95% O2 and 5% CO2). Each slice was transferred to a submerged chamber mounted on an upright microscope and neurons were visualized using differential interference contrast infrared videomicroscopy. For the recordings, patch pipettes (5 to 10 megohm) were pulled from borosilicate glass and filled with an intracellular solution containing: 135 K-gluconate, 4 NaCl, 10 HEPES, 2 MgCl2, 0.5 EGTA, 2.5 Na2-ATP, 0.5 Na2-GTP, pH adjusted to 7.4, 290-300 mOsmol. Recordings were performed with MultiClamp-700B (Molecular Devices) at 35°. The excitability curves were performed by holding the membrane resting potential at -65 mV and injecting 2s-positive current steps from 0 to 490 pA. The membrane resistance and membrane capacitance were determined by repeated injections of a -20pA current step. The liquid junction potential was not compensated.

#### Immunohistochemistry

After patch-clamp recordings, slices were fixed in Antigenfix (Diapath, P0014) for 1h and washed five times for 3 minutes in Tris-buffered saline (TBS). Immunodetection was done in free-floating sections. Slices were treated with 50 mM NH4Cl for 30 min and incubated in blocking buffer (10% donkey serum, 0.3% Triton X-100, TBS/BSA 0.25%) for 4 h to avoid nonspecific binding. After washing in TBS/Triton X-100 0.3% five times for 5 min, slices were incubated for 48h at 4 °C with the primary antibodies diluted in incubation buffer (TBS/BSA 0.25%, Triton X-100 0.3%, 1% donkey serum). The primary antibodies used were: rabbit anti-CamKIIα (1:200, Proteintech, 13730-1-AP) and mouse anti-ankyrinG (1:100, Sigma-Aldrich, MABN466). After washing in TBS/Triton X-100 0.3% three times for five minutes, the secondary antibodies, Cy3 donkey anti-rabbit (1:200, Jackson ImmunoResearch, 711-165-152) and Cy2 donkey anti-mouse (1:200, Jackson ImmunoResearch, 715-545-151) were incubated for 2 h at room temperature and then washed in TBS/Triton 0.3 %, six times for 3 minutes. After staining, the slices were mounted with Fluoromount-G (Invitrogen 00-4959-52). The images were acquired on Apotome 3 Zeiss microscope with a 20x objective. AIS length and distance from the soma were analyzed using the SNT plugin ImageJ/Fiji software by manual detection. Soma surface were analyzed using the Polygon function of ImageJ/Fiji software by manual detection. All analysis were performed blindly to treatment and sex.

#### Hodgkin-Huxley modelling

Computer simulations for Hodgkin-Huxley models were done in NEURON 8.2.6 software, all the codes are available at https://github.com/mizbili/Axon-initial-segment-and-facial-neuropathy. We used a ball-and-stick morphology composed of one soma, one hillock and one AIS. Simulations were run with 0.1-ms time steps, and the nominal temperature of simulation was 34°C. Soma and hillock contained passive channels, Ih channels, one Nav type (Nav1.2), 2 Kv types (K_Tst and SKv3_1), one calcium-activated potassium channels type (SK_E2), one Cav type (Ca_HVA) and one mechanism of intracellular Ca^2+^ (CaDynamics_E2). AIS contained passive channels, two Nav type (Nav1.2 and Nav1.6), 1 Kv type (SKv3_1), one calcium-activated potassium channels type (SK_E2), one Cav type (Ca_HVA) and one mechanism of intracellular Ca^2+^ (CaDynamics_E2). All channels were taken from the pool released by the Blue Brain Project^4,5^ except for Nav1.2 and Nav1.6 that were adapted to follow the biophysics of sodium channels recorded in AIS of layer 5 pyramidal neurons^6^. Electrophysiological parameters presented a uniform distribution in each compartment except for Nav1.2 which presented a sigmoidal decrease along AIS and Nav1.6 a sigmoidal increase along AIS^6^. In fact, distribution of Nav1.2 and Nav1.6 conductances followed the equations:

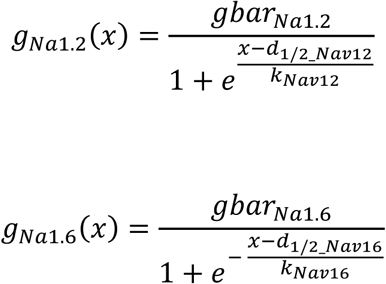

With x the distance from the AIS beginning, gbar_Na12_ and gbar_Na16_ the maximal conductances, d_1/2_Na12_ and d_1/2_Na16_ the midpoints of the sigmoidal curves, k _Na12_ and k _Na16_ the factors determining the slope of the sigmoidal curves. Optimization of the different parameters was done using BluePyOpt software^7^ (see Supplementary Table 1 for the values of the parameters). For the male model, the IONL condition was simulated simply by increasing AIS length without soma-AIS distance modification (Supplementary Table 2), as seen experimentally. As we observed experimentally that somas were smaller in female than in male, the female model was obtained by a simple soma size decrease of the male model (Supplementary Table 2) without modification of the electrophysiological parameters. For the female model, the IONL condition was simulated by increasing AIS length and decreasing soma-AIS distance (Supplementary Table 2), as seen experimentally. The stimulations were done, as in experimentally measured I/F curves, by 2-s long current steps ranging from 0 to 300 pA.

#### Network modelling

The network modelling was performed using python self-written codes available at https://github.com/mizbili/Axon-initial-segment-and-facial-neuropathy.

#### Single neuron models

The single neurons models composing the networks were built using monocompartmental conductance-based adaptative leaky integrate-and-fire (CAd-LIF) model^8–10^ using the equations:

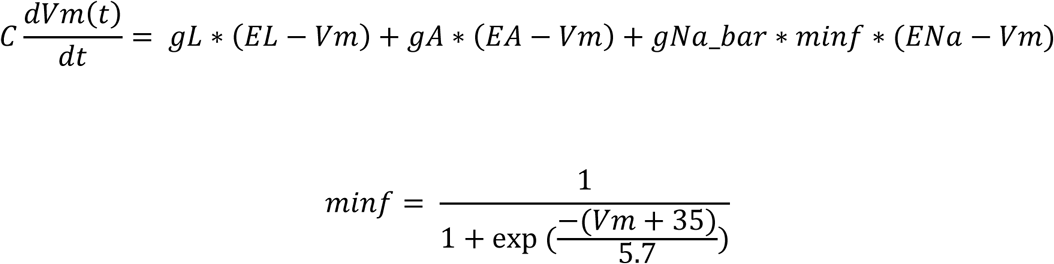

with C is the membrane capacitance, gL the leak conductance, EL the reversal potential of leak conductance, gA the subthreshold adaptation conductance, EA the reversal potential of adaptation conductance, gNa_bar the maximal sodium conductance, minf the activation curve of sodium conductance, ENa the reversal potential of sodium conductance. The adaptation conductance obeyed to:

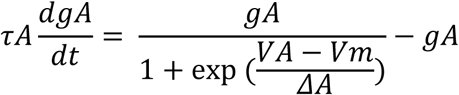

with τA the time constant of adaptation, gA_bar the maximal subthreshold adaptation conductance, VA is the subthreshold adaptation activation potential, and ΔA is the slope of subthreshold adaptation. When Vm crossed a threshold V_AP_, it was reset to a reset voltage VR and gA was incremented using the following equations:

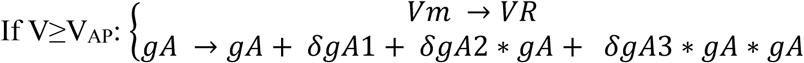

The different parameters were determined to fit the excitability curves of Sham and IONL neurons recorded in female and male as well as excitability curves of fast-spiker neurons recorded in Sham and IONL animals recorded in both sexes. The common parameters to all the models are: V_AP_ = -20 mV, EA = -80.07 mV, EL = -78 mV, ENa = 50 mV, VA = -36.89 mV, ΔA = 5.58 mV. The other parameters are described in Supplementary Table 3.

#### Networks models

Using the monocompartmental single neurons models, we built 4 networks types: a Sham-like female network, a IONL-like female network, a Sham-like male network and a IONL-like male network.

A given network was composed of 800 pyramidal-like neurons (which properties depending on the network) and 200 FS-like interneurons. The connection probability between two neurons was 10%. To simulate the synapses, we used the equations described in Zbili et al., 2020^11^. Each presynaptic spike produces an exponentially decaying current with time constant τexc = 5 ms for excitatory synapses and τinh = 10 ms for inhibitory synapses. A given synapse followed the equation:

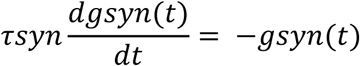

where syn ∈ {exc, inh} and each presynaptic spike trigger an instantaneous increase: gsyn → gsyn + λsyn*wi, where wi is the synaptic weight and λsyn is a scaling factor calculated so that a synaptic weight of 1 mV produces PSPs of peak size 1 mV. Note that gsyn is in units of volt, i.e., the membrane resistance is implicitly included in the variable. For exponential synapses, we can compute analytically

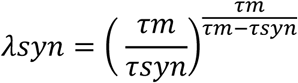

where τm is the membrane time constant of the postsynaptic neuron given by the equation:

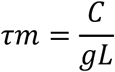

For excitatory synapses, the synaptic weights wi were randomly chosen using a lognormal law of mean 0.2 mV and sd 1 mV. For inhibitory synapses, the synaptic weights wi were randomly chosen using a normal law of mean -30 mV and sd 4 mV. Finally, to simulate external inputs, the pyramidal-like cells were stimulated using random excitatory synapses trains at various frequencies (from 0.2 to 60 Hz) to test the influence of external inputs on the network activity. For each condition tested (Sham-like female, Sham-like male, IONL-like female, IONL-like male) we ran 20 trials per external input frequency. At each trial, the connection matrix, the synaptic weights and the external inputs trains were randomly chosen as described above.

#### Statistics

Statistical analyses were performed using mixed linear model regressions (MLMR) to account for both fixed and random sources of variability^12^. Pairwise t-tests p-values were then obtained from the marginal means and the variance derived from the models. P-values for multiple tests were corrected by Benjamini-Hochberg False Discovery Rate method. Model residuals were inspected using quantile–quantile (QQ) plots of residuals to verify assumption of normality. When this assumption was violated, we performed a Box-Cox transformation^13^ to normalize the data and re-applied the MLMR to the transformed data (this transformation was necessary for rheobase, gain, AP half-width, AIS distance and Soma area). To observe the variations of parameters in function of the day post-surgery, we used linear regressions. All the analyses were made using statsmodels and scipy python libraries. The codes are available at https://github.com/mizbili/Axon-initial-segment-and-facial-neuropathy.

---

**Supplementary Figure 1.**
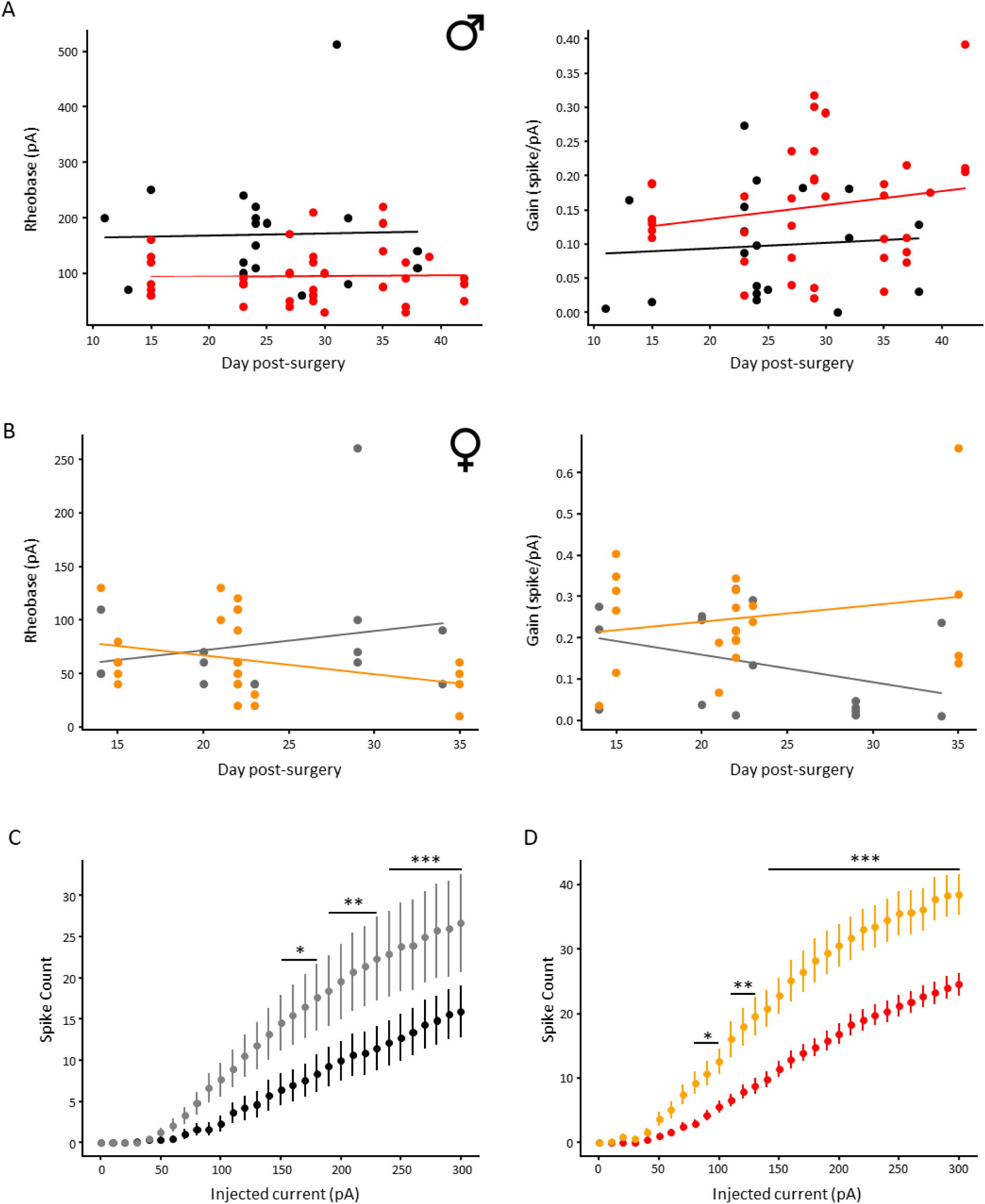
Analyses of Input/Output curves with the day post-surgery and between sexes. **(A)** No correlation of rheobase or gain with day post-surgery in Sham (black) or IONL (red) male rats. **(B)** No correlation of rheobase or gain with day post-surgery in Sham (grey) or IONL (orange) female rats. **(C)** Stronger L5PC excitability in females (grey) than males (black) in Sham condition. **(D)** Stronger L5PC excitability in females (orange) than males (red) in IONL condition.

**Supplementary Figure 2.**
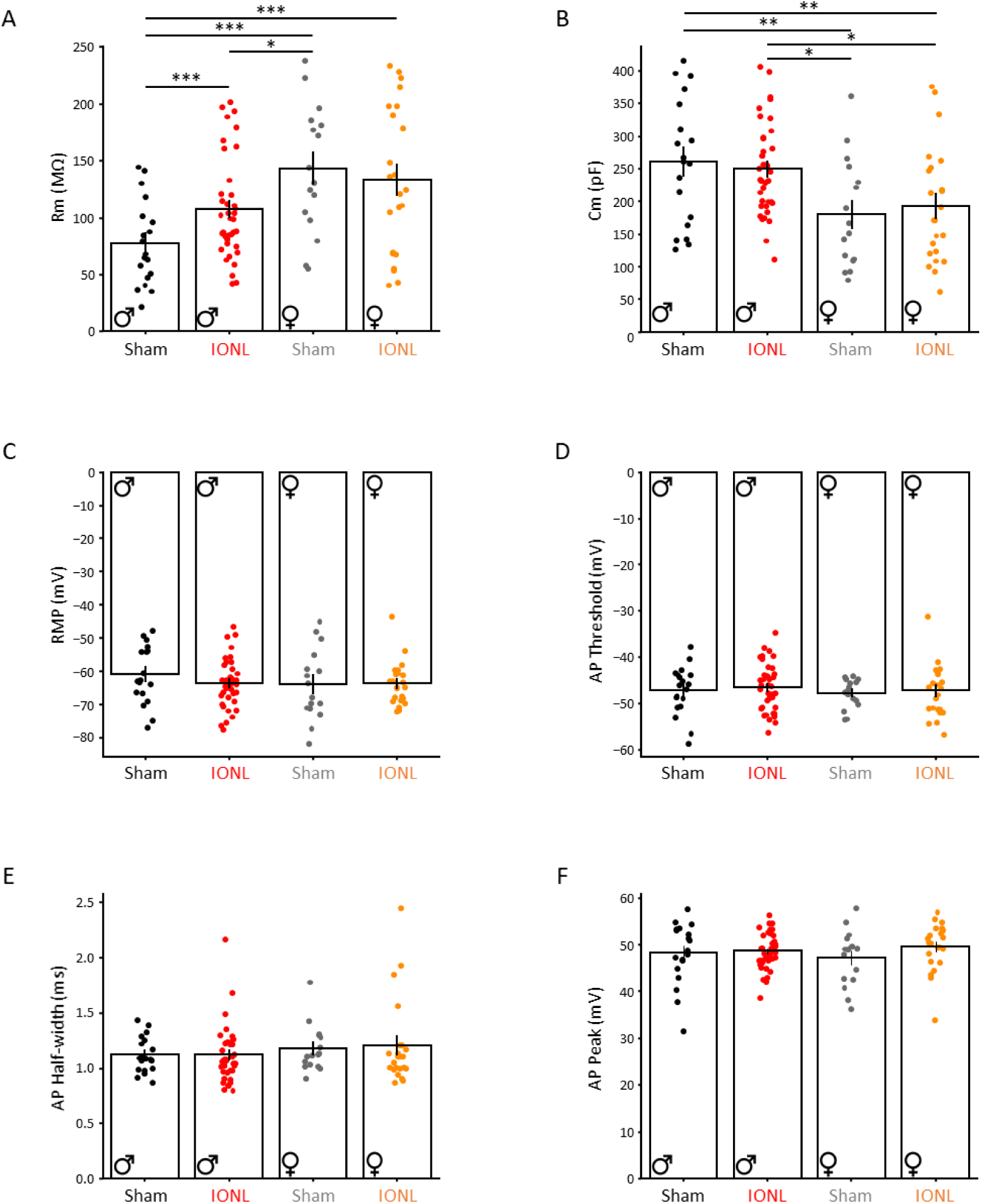
Electrophysiological parameters in Sham and IONL rats. **(A)** L5PC membrane resistance in Sham and IONL male and female rats. Note that Rm is increased following IONL in male but not in female rats. Note that female L5PC display larger Rm than male in Sham animals, suggesting a sexual dimorphism of this parameter for this neuronal type. **(B)** L5PC membrane capacitance in Sham and IONL male and female rats. Note that Cm is not modified by IONL in male nor in female rats. Note that female L5PC display smaller Cm than male L5PC, suggesting a sexual dimorphism of this parameter for this neuronal type. **(C)** No difference in L5PC resting membrane potential in Sham and IONL male and female rats. **(D)** No difference in L5PC AP threshold in Sham and IONL male and female rats. **(E)** No difference in L5PC AP half-width in Sham and IONL male and female rats. **(F)** No difference in L5PC AP peak in Sham and IONL male and female rats.

**Supplementary Figure 3.**
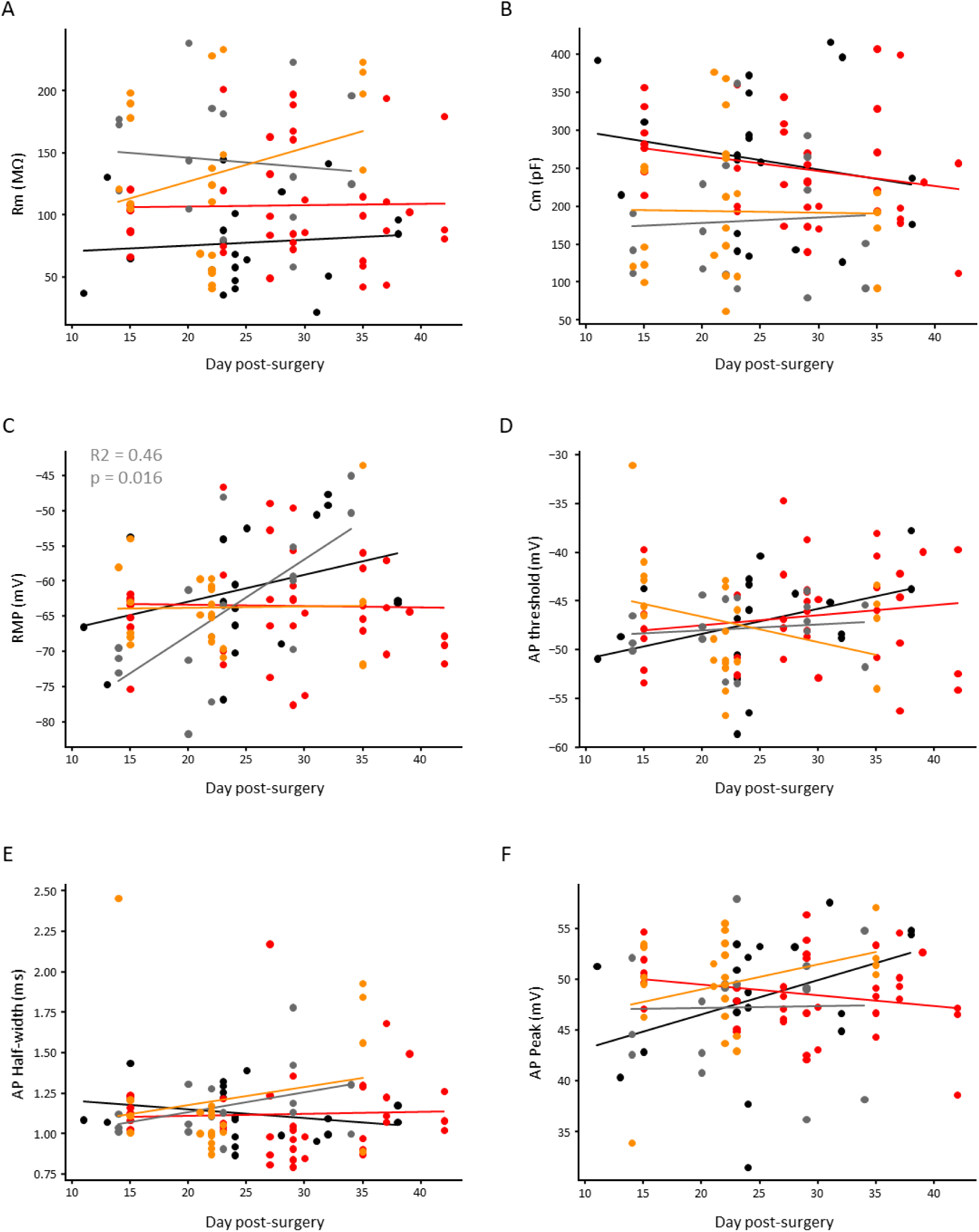
No correlation of electrophysiological parameters with the day post-surgery in the recorded neurons. (A-F) Correlation tests of Rm, Cm, RMP, AP threshold, AP half-width and AP peak with day post-surgery in Sham male (black), IONL male (red), Sham female (grey) and IONL female (orange) rats. No correlation was found except for a RMP depolarization with day post-surgery in Sham female rats. R^2^ and p-values of statiscally significant tests are displayed on the graphs.

**Supplementary Figure 4.**
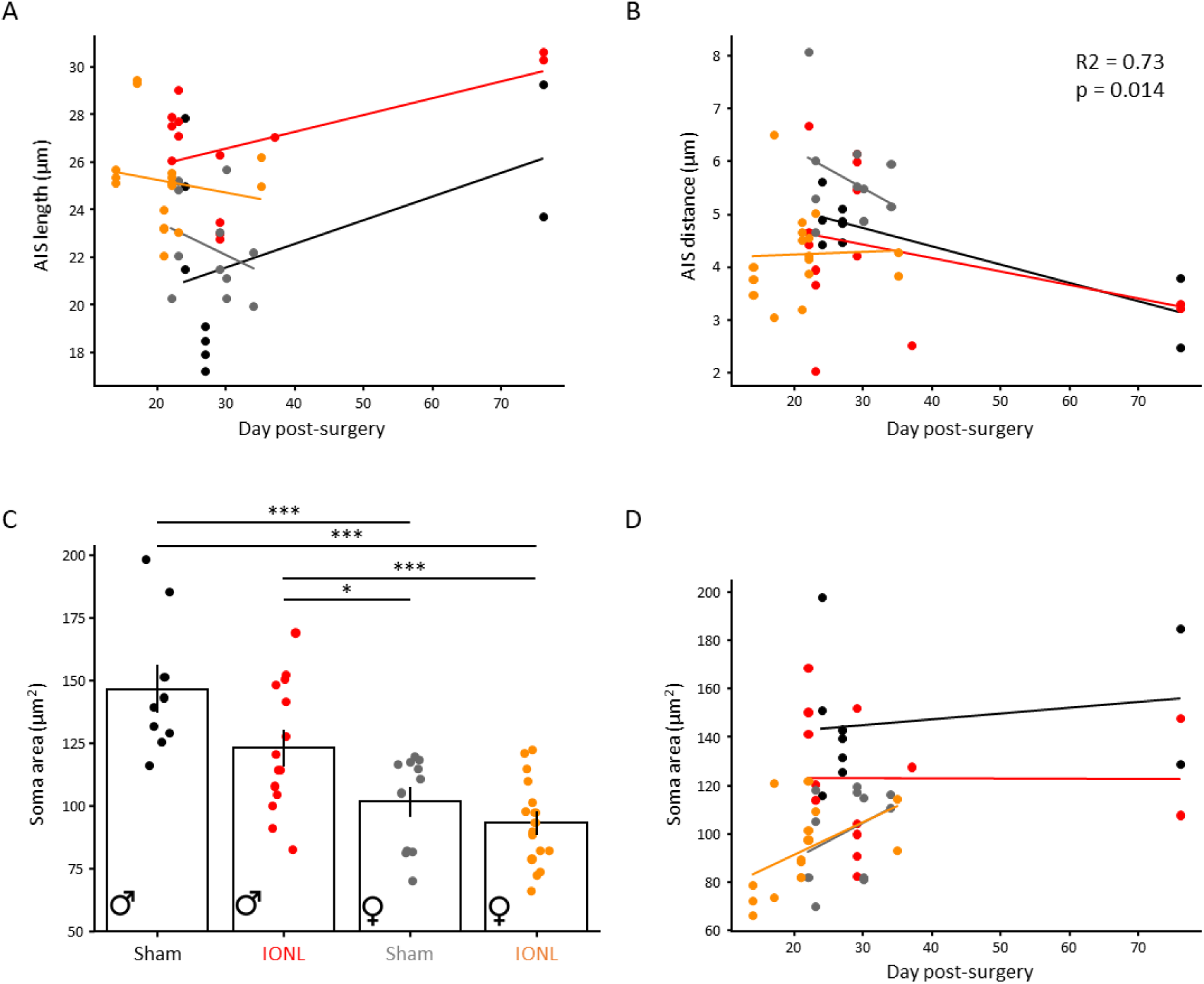
Analyses of axon initial segment and soma area. **(A)** Correlation tests of AIS length with day post-surgery in Sham male (black), IONL male (red), Sham female (grey) and IONL female (orange) rats. No correlation was found. **(B)** Correlation tests of soma-AIS distance with day post-surgery in Sham male (black), IONL male (red), Sham female (grey) and IONL female (orange) rats. No correlation was found except for a reduction of the distance in Sham male (R^2^ and p-values of statiscally significant tests are displayed on the graphs). **(C)** L5PC soma area in Sham and IONL male and female rats (Sham males: 781 soma from 9 slices in 4 animals, IONL males: 1280 soma from 13 slices in 6 animals; Sham females: 1025 soma from 11 slices in 5 animals, IONL females: 2725 soma from 16 slices in 6 animals). Note that IONL did not induce modification of soma area in males nor in females while female displayed smaller soma area males (each dot represents the mean of slice). **(D)** Correlation tests of soma area with day post-surgery in Sham male (black), IONL male (red), Sham female (grey) and IONL female (orange) rats. No correlation was found.

**Supplementary Figure 5.**
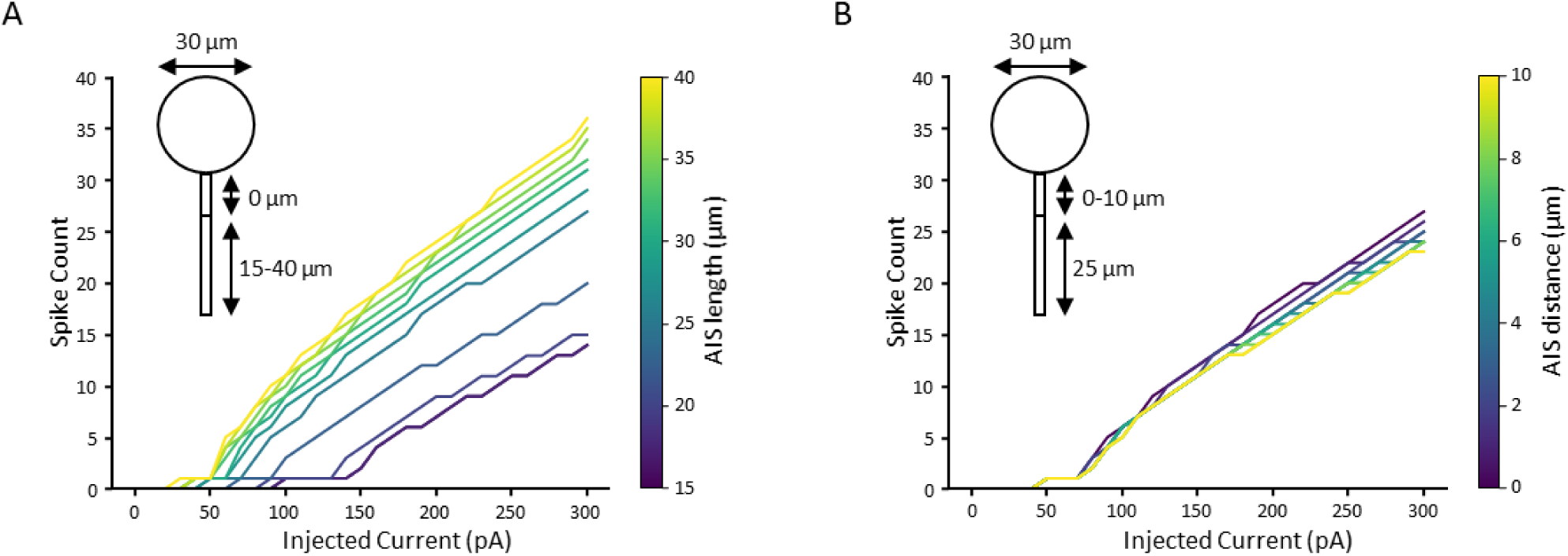
Stronger effect of AIS length than soma-AIS distance on model excitability. **(A)** AIS length increase from 15 to 40 µm induce large enhancement of model excitability. **(B)** Soma-AIS distance increase from 0 to 10 µm induce small decrease of model excitability.

**Supplementary Figure 6.**
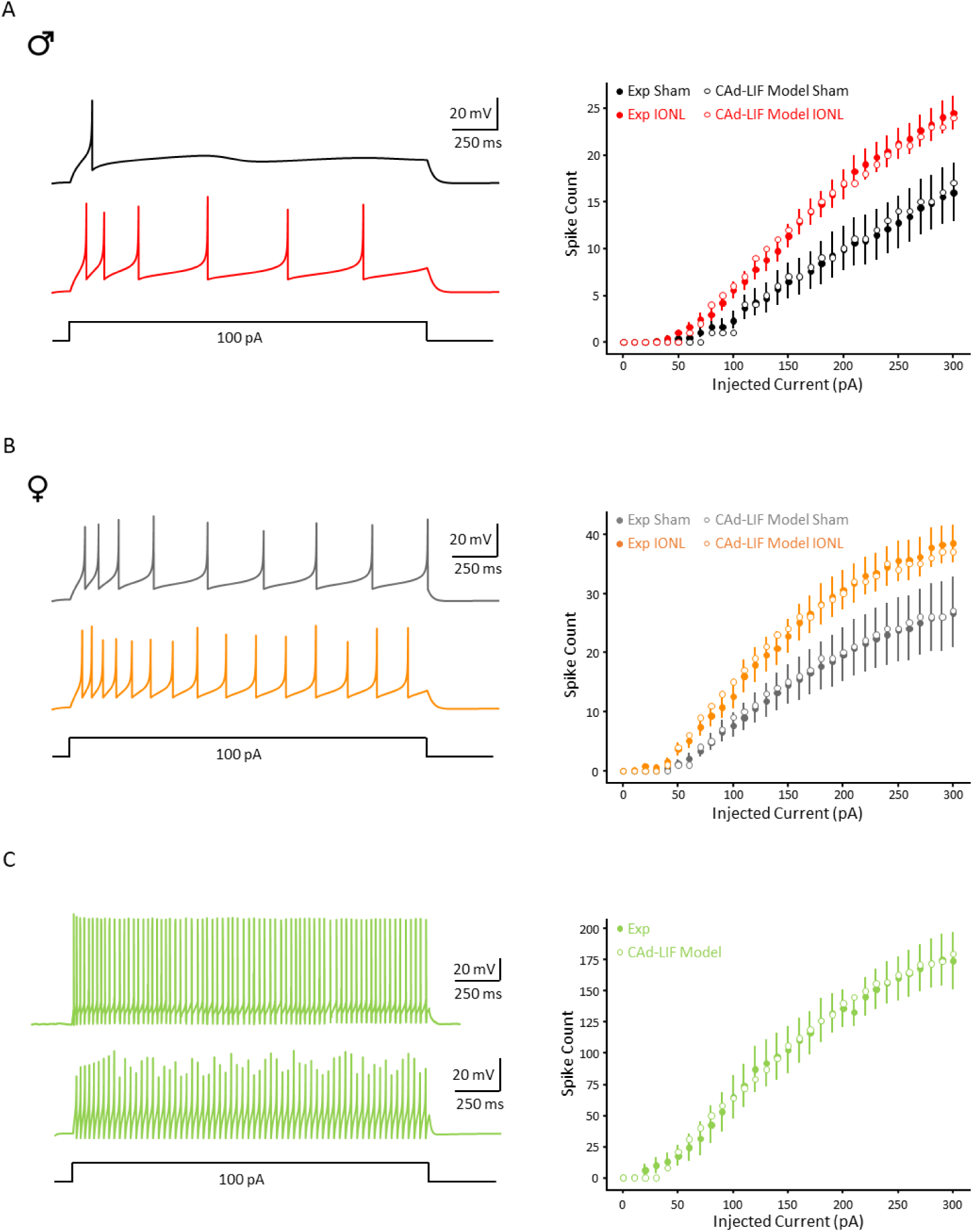
Input/Output curves of CAd-LIF single neurons used for network models. **(A)** Left, Current-clamp traces obtained by injection of a 100 pA-current step in the Sham male model (black) and the IONL male model (red). Right, Experimental (full dots) and modeled (empty dots) I/F curves in Sham male (black) and IONL male (red) conditions. **(B)** Left, Current-clamp traces obtained by injection of a 100 pA-current step in the Sham female model (grey) and the IONL female model (orange). Right, Experimental (full dots) and modeled (empty dots) I/F curves in Sham female (grey) and IONL female (orange) conditions. **(C)** Up Left, Example of a current-clamp FS interneuron recording. Low Left, Current-clamp trace obtained by injection of a 100 pA-current step in the FS interneurons model. Right, Experimental (full dots) and modeled (empty dots) I/F curves of Fast-Spiker inhibitory neurons.

**Supplementary Table S1.** Electrophysiological parameters for the Hodgkin-Huxley model.

| Parameter | Compartment |  |
| --- | --- | --- |
|  | Soma/Hillock | AIS |
| cm ($\mu\text{F}/\text{cm}^2$) | 4.975549185 | 4.975549185 |
| Ra ($\Omega\text{cm}$) | 100 | 100 |
| gpas ($\text{S}/\text{cm}^2$) | 1.57E-05 | 1.57E-05 |
| gbarlh ($\text{S}/\text{cm}^2$) | 8.30E-05 | |
| gbarK_Tst ($\text{S}/\text{cm}^2$) | 0.029162317 | |
| gbarSKv3_I ($\text{S}/\text{cm}^2$) | 0.033231362 | 6.945241481 |
| gbarSK_E2 ($\text{S}/\text{cm}^2$) | 0.001645799 | 0.001800827 |
| gbarCa_HVA ($\text{S}/\text{cm}^2$) | 4.63E-05 | 1.35E-06 |
| gamma_CaDynamics_E2 | 0.006897399 | 0.009211661 |
| decay_CaDynamics_E2 (ms) | 321.0571592 | 295.4420659 |
| minCai_CaDynamics_E2 (mM) | 0.00026639 | 1.33E-04 |
| gbarNav12 ($\text{S}/\text{cm}^2$) | 1.001871288 | 6.718166475 |
| gbarNav16 ($\text{S}/\text{cm}^2$) | | 17.17086288 |
| d1/2_Nav12 ($\mu\text{m}$) | | 28.49935279 |
| kNav12 ($\mu\text{m}^{-1}$) | | 3.482709277 |
| d1/2_Nav16 ($\mu\text{m}$) | | 20.42669878 |
| kNav16 ($\mu\text{m}^{-1}$) | | 1.335146383 |
| epas (mV) | -69.78774055 | -69.78774055 |
| ena (mV) | 50 | 50 |
| ek (mV) | -85 | -85 |
| eca (mV) | 100 | 100 |
| eih | -45 |  |

**Supplementary Table S2.** Morphological parameters for the Hodgkin-Huxley model.

| Parameter | Model Condition |  |  |  |
| --- | --- | --- | --- | --- |
|  | Male Sham | Male IONL | Female Sham | Female IONL |
| Soma L ($\mu\text{m}$) | 30 | 30 | 24.40230545 | 24.40230545 |
| Soma diam ($\mu\text{m}$) | 30 | 30 | 24.40230545 | 24.40230545 |
| Hillock L ($\mu\text{m}$) (soma-AIS distance) | 4.4 | 4.4 | 5.6 | 4.4 |
| Hillock diam ($\mu\text{m}$) | 1 | 1 | 1 | 1 |
| AIS L ($\mu\text{m}$) | 22.3 | 26 | 22.3 | 26 |
| AIS diam ($\mu\text{m}$) | 1 | 1 | 1 | 1 |

**Supplementary Table S3.** Parameters for the CAd-LIF models used in network modelling.

| Models | Parameter |  |  |  |  |  |  |  |  |
| --- | --- | --- | --- | --- | --- | --- | --- | --- | --- |
| | C (pF) | gL (nS) | gNA_bar (nS) | gA_bar (nS) | VR (mV) | $\delta gA1$ (nS) | $\delta gA2$ (nS) | $\delta gA3$ (nS) | $\tau A$ (ms) |
| male sham pyramidal neuron | 282.74 | 8.53 | 84.82 | 282.74 | -70 | 1.1 | 0.031 | 0 | 506.62 |
| female sham pyramidal neuron | 282.74 | 8.53 | 124.41 | 282.74 | -70 | 0.3 | 0.003 | 0.006 | 506.62 |
| male IONL pyramidal neuron | 282.74 | 8.53 | 113.1 | 282.74 | -70 | 0.4 | 0.073 | 0 | 506.62 |
| female IONL pyramidal neuron | 282.74 | 8.53 | 155.51 | 282.74 | -70 | 0.07 | 0.001 | 0.0052 | 506.62 |
| FS interneuron | 70.69 | 7.07 | 106.03 | 70.69 | -80 | 0.05 | 0.014 | 0.004 | 150 |

